# A SNAI2-PEAK1 stromal axis drives progression and lapatinib resistance in HER2-positive breast cancer by supporting a cytokine expression profile that converges on PI3K/Akt signaling

**DOI:** 10.1101/2020.05.15.098772

**Authors:** Sarkis Hamalian, Robert Güth, Farhana Runa, Justin Molnar, Eric Vickers, Megan Agajanian, Jonathan Humphries, Martin J. Humphries, Julia Tchou, Ioannis K. Zervantonakis, Jonathan A. Kelber

**Author notes:** Current address: Department of Bioengineering, University of Pittsburgh, Center for Bioengineering, 300 Technology Dr., Pittsburgh, PA 15219.

## Abstract

Intercellular mechanisms by which the stromal microenvironment contributes to solid tumor progression and targeted therapy resistance remain poorly understood, presenting significant clinical hurdles. PEAK1 (Pseudopodium-Enriched Atypical Kinase One) is an actin cytoskeleton- and focal adhesion-associated pseudokinase that promotes cell state plasticity and cancer metastasis by mediating growth factor-integrin signaling crosstalk. Here, we determined that stromal PEAK1 expression predicts poor outcomes in HER2-positive breast cancers high in SNAI2 expression and enriched for MSC content. Notably, we identified that mesenchymal stem cells (MSCs) and cancer-associated fibroblasts (CAFs) express high PEAK1 protein levels and MSCs require PEAK1 to potentiate tumorigenesis, lapatinib resistance and metastasis of HER2-positive breast cancer cells. Analysis of PEAK1-dependent secreted factors from MSCs revealed a CCL4-, INHBA- and GDF5-focused network that converged on PI3K/Akt signaling. In this regard, we observed that MSC expression of PEAK1 is required for sustained Akt phosphorylation in neighboring HER2-positive breast cancer cells following lapatinib treatment. Finally, we uncovered a significant correlation between INHBA and PEAK1 expression levels in breast cancer, and that INHBA is an excellent predictor of disease relapse and decreased survival in HER2-positive tumors enriched for PEAK1 expression and MSC content. Importantly, we provide the first evidence that PEAK1 promotes tumorigenic phenotypes through a previously unrecognized SNAI2-PEAK1-INHBA-PI3K/Akt stromal to tumor cell signaling axis. These results establish a new, targetable intercellular mechanism that may be leveraged to improve targeted therapy responses and patient outcomes in breast cancer and other stroma-rich malignancies.

## INTRODUCTION

Cell state plasticity enhances intratumoral heterogeneity and has been shown to be a culprit underlying metastasis, therapy resistance and progression in cancer [1–3]. Previous studies have demonstrated a causative relationship between increased stromal tissue content (i.e., desmoplasia), including cancer associated fibroblasts (CAFs) or mesenchymal stem cells (MSCs), in breast cancers and lapatinib resistance or metastasis [4–7]. In the case of HER2-positive breast cancer, where upregulation of the receptor tyrosine kinase HER2 (ErbB2) occurs in approximately 20% of all breast malignancies [8], both trastuzumab- and lapatinib-based regimens offer significant clinical benefit [9]. However, a substantial percentage of these tumors display either primary resistance or may be initially sensitive but then adapt to develop acquired resistance [10], and clinical work suggests that patients who progress on lapatinib therapy commonly develop metastatic disease [11]. Importantly, stromal cell non-autonomous mechanisms by which the tumor microenvironment drives lapatinib resistance and/or resistance-associated metastasis remain poorly understood.

Pseudopodium-Enriched Atypical Kinase One (PEAK1 or SGK269) is a cytoskeleton-associated pseudokinase [12] and member of the new NFK3 kinase family that has been demonstrated to play key roles in cancer initiation and progression across multiple cancer types including breast [13–15], pancreatic [12, 16], lung [17] and colon [12, 18, 19]. Importantly, two other NFK3 kinase family members (SGK223 or Pragmin and PEAK3) have been recently demonstrated to also regulate cancer progression [20, 21]. We previously characterized breast cancer cell autonomous functions of PEAK1 and upstream eIF5A1/2-dependent translation in mediating epithelial-mesenchymal transition (EMT), metastasis and transforming growth factor beta (TGFβ)/fibronectin signaling [13–15, 22]. In this regard, PEAK1 has been previously identified as part of the meta-adhesome [23] and a regulator of focal adhesion dynamics [24]. Notably, PEAK1 was recently determined to be a core constituent of the fibroblast adhesome using multiplexed proximity biotinylation methods [25]. In further support of a role for PEAK1 in mediating growth factor-integrin crosstalk, Zheng and colleagues reported that PEAK1 is a critical adaptor protein govering Shc1 association with cytoskeletal reorganization, trafficking and signal termination proteins downstream of EGF/Akt/PTPN12 activity to mediate cell invasion [26]. Still, in the context of tumorigenesis, metastasis and/or therapy resistance, the function and mechanism of action for PEAK1 within the stromal microenvironment remain unknown.

Given that PEAK1 associates with the actin cytoskeleton and focal adhesion complexes in fibroblast cell types [12, 23–25], and fibroblasts have well-documented roles within the cancer stroma, we addressed whether PEAK1 may promote tumorigenesis via the non-epithelial compartment of solid tumors. To this end, we report that PEAK1 expression in breast cancer stroma is associated with relapse in HER2-positive breast cancer and that PEAK1 is predominantly expressed in SNAI2-positive fibroblast-like cells within the tumor mass. In agreement with these data, patient-derived CAFs and MSCs express high levels of PEAK1 and can promote malignant phenotypes and lapatinib resistance *in vitro* and *in vivo* in a PEAK1-dependent manner. Finally, we highlight a previously unrecognized PEAK1-INHBA-Akt stromal to tumor cell signaling axis that may be targeted to abrogate therapeutic resistance in HER2-positive breast cancer and improve patient outcomes.

## MATERIALS AND METHODS

### Cell Culture

RAW264.7, C3H10T1/2, NIH3T3, Swiss3T3, EA.hy926, BT474 and MCF7 cells were purchased from the American Tissue Culture Collection (ATCC). Patient-derived cancer-associated fibroblasts (CAFs) (i.e., TB123, TB125, TB98, TB130, TB122, TB129) were provided by Dr. Julia Tchou’s laboratory of the Perleman Center for Advance Medicine at the University of Pennsylvania. Isolation and maintenance protocols of these CAFs have been described [27]. Py230 cells were provided by Dr. Lesley Ellies’ laboratory at University of California San Diego. BT474-H2BeGFP and AR22 cells were provided by Dr. Joan Brugge’s laboratory at the Ludwig Center for Cancer Research & Department of Cell Biology at Harvard Medical School. RAW264.7, C3H10T1/2, NIH3T3, Swiss3T3, EA.hy926 and AR22 cells were cultured in Dulbecco’s Modified Eagle’s Medium (DMEM)/High glucose growth media supplemented with 10% fetal bovine serum (FBS), 1% penicillin/streptomycin and 0.1% gentamycin. MCF7 cells were cultured in DMEM High glucose growth media supplemented with 10% fetal bovine serum (FBS), 1% penicillin/streptomycin and 0.1% gentamycin and 0.01 mg/ml human recombinant insulin. BT474 cells were cultured in Rosewell Park Memorial Institute (RPMI-1640) growth media supplemented with 10% FBS, 1% penicillin/streptomycin and 0.1% gentamycin. CAF lines were cultured in (DMEM)/High glucose growth media supplemented with 20% fetal bovine serum (FBS), 1% penicillin/streptomycin and 0.1% gentamycin. Py230 cells were cultured in F-12K growth media supplemented with 5% fetal clone serum (FCS), 1% penicillin/streptomycin, 0.1% gentamycin and 0.1% MITO+ Serum Extender. Cultures were maintained at 37°C with 5% CO_2_.

### Bioinformatics

Target gene/protein transcript levels in normal and breast cancer stroma or total breast cancer tissues were extracted from microarray or sequencing data published by Karnoub et al. 2006 Cell, Finak et al. 2008 Nat. Med., Pereira et al. 2012 Nat., Nagy et al. 2018 Sci. Rep. and/or Gyorffy et al. 2010 Breast Cancer Res Treat and deposited in the Oncomine, Cancer BioPortal, or KMPlot databases. Similarity matrices were generated using the Morpheus website. Immunohistochemical protein data from patient breast tumors are available in the Human Protein Atlas (http://www.proteinatlas.org). Interactome networks and gene set enrichments (GSEs) were generated using the Core Expression Analysis application within the Ingenuity Pathway Analysis (IPA) platform.

### Conditioned Media (CM)

Cells were plated in 10 cm plates at 6e5 cells and incubated until 70% confluent. Media in each 10 cm plate was changed to 6 mL of appropriate media without serum and incubated for an additional 48 hours. Mock/control media was made by placing the same media into a plate without cells for 48 hours. Media was collected, centrifuged at 1000 rpm for 5 minutes and used right away or stored at −80°C in 15 mL aliquots until needed. Before the CM was used in experiments, it was diluted 1:1 with the appropriate fresh serum-free media.

### Western Blot

Whole cell extracts (i.e., cell lysates) were collected in standard radioimmunoprecipitation assay (RIPA) buffer containing phosphatase and protease inhibitors and rotated at 4°C overnight before pelleting insoluble material and protein analysis of supernatants. Lysate protein concentrations were determined by Bradford assay or bicinchoninic acid (BCA) assay. Proteins in lysates, normalized for 20 ug of protein/well, were reduced in 4× LDS sample buffer with DTT and separated using 4-12% Bis-Tris NuPage gels. Gels were blotted onto nitrocellulose membranes and probed for each antigen using the following antibodies at the indicated dilutions: Human PEAK1 (Millipore 1:400), Mouse PEAK1 (Abgent 1:400), α-tubulin (Pro-Sci 1:1000), GAPDH (Pro-Sci 1:1000) and β–actin (Pro-Sci 1:1000). Secondary antibodies were used at a 1:5000-1:10,000 dilution. Developed autoradiography films were scanned and analyzed for relative band intensities using Fiji software after image thresholding.

### Immunofluorescence

Indicated cells fixed using 4% paraformaldehyde for 20 minutes, permeabilized for 10 minutes in 0.1% Triton-X100, blocked in 10% BSA and stained with the indicated primary/secondary antibodies at 1:400 in 2% BSA in PBS. Images were collected using a Leica DMI6000 inverted microscope at 40X magnification.

### Chorioallantoic Membrane Assay (CAM Assay)

Rhode Island Red Hatching Eggs were purchased from Meyer Hatchery and incubated at 38 degrees Celsius and 60% humidity for 10 days on an automated egg turner. On day 10, the shell was sterilized with Rocadyne (RMC) and a small hole was made at the blunt end of the egg to evacuate the air pocket and pull the CAM away from the shell and shell membrane. While candeling the egg, a small rectangular window was cut over the umbilical vein using a dremel (Black and Decker). A sterile silicon ring was placed onto the CAM in an area of dense vasculature. Cells were trypsinized, counted and resuspended in ice-cold 100% GFR Matrigel using ice-cold pipette tips at the following specifications: mono-xenografts had 1e6 cells suspended in 40μl of Matrigel per embryo and co-xenografts of breast cancer cells with stromal cells had 1e6 cells per line suspended in 40μl of Matrigel per embryo. Xenografts and co-xenograft suspensions were then applied to the CAM using ice-cold pipette tips. Sterile surgical tape was applied to the windowed area as well as the blunt end and the eggs were placed back into the incubator without egg turning. In experiments with drug treatments, two days after xenografting, the eggs were re-sterilized and the surgical tape was carefully peeled back to reveal one corner of the window. Drugs are prepared at 2× the final concentration (i.e., 3600 nM or 2 μM lapatinib) in 40 uL of ice-cold 100% GFR Matrigel per embryo. At day 17, the embryo was then extracted from the egg and the developing chicken sacrificed via decapitation. The primary CAM tumor was dissected, brain and lung tissues were weighed and divided into two samples each for flash-freezing (in an EtOH/dry ice bath) and fixing (in 10% Formalin), respectively.

### Quantitative Polymerase Chain Reaction (qPCR)

20mg of relevant tissues were weighed, homogenized using a micro-homegenizer (ClaremontBio Solutions) in digestion solution and then heated at 56°C for 3.5 hours. Genomic DNA was then purified using GeneJet Genomic DNA purification kit (ThermoFisher Scientific) and the concentration measured by NanoDrop. gDNA was stored at −20°C for downstream qPCR analysis. Samples of gDNA were diluted with nuclease free water so that addition of 11.25 μl of this diluted sample delivers 56.25 ng of gDNA. Primers were purchased from Integrated DNA for human Alu repeat (sense: 5′ ACG CCT GTA ATC CCA GCA CTT 3′ and antisense: 5′ TCG CCC AGG CTG GAG TGC A 3′) and chicken gapdh (sense: 5′ GAG GAA AGG TCG CCT GGT GGA TCG; antisense: 5′ GGT GAG GAC AAG CAG TGA GGA ACG) and used at a concentration of 10nmol/mL per reaction. A master mix was made so that each well would contain 12.5 μL of Maxima SYBR® Green (Thermo Scientific) and 1.25 μLl of gene-specific primer. Contents for experimental wells include 11.25 μL diluted gDNA and 13.75 μL of master mix. qPCR plates were processed using the ABI7300 instrument with the following thermal cycles settings: Stage 1, 1 repetition - 50°C for 2 minutes / Stage 2, 1 repetition - 95°C for 30 seconds / Stage 3, 40 repetitions at 95°C for 15 seconds / Stage 4 at 62°C for 1 minute. Relative metastasis was calculated as previously described [28]. Briefly, dCt values were calculated as aluCt - gapdhCt. ddCt values were calculated as dCt_variable_-Average dCt_control_. RQ values were calculated as 2e^(−ddCt)^.

### Lentiviral Transduction

Cells were plated at 4.8e6 cells/well into a 6 well plate and left to attach overnight. Viral mixes were created with an aliquot of virus into complete media and polybrene at 8μg/ml (Sigma-Aldrich) to have a target multiplicity of infection (MOI) of 5. Viral particles contained a puromycin resistant pKLO.1 vector with a scramble shRNA or PEAK1-specific shRNA (5 different constructs). Viral mixes were added to their respective wells and left to incubate for 24 hours, after which regular media was replaced. The following day, media was changed and supplemented with 1 ug/mL puromycin. Cells were expanded and knockdown efficacy was validated by Western blotting.

### Immunohistochemistry

Tissue samples were sent to the UCLA Tissue Procurement Core Laboratory for paraffin embedding, tissue sectioning and H&E staining. The tissue was dehydrated and incubated in primary antibodies (αSMA, 1:200) overnight. Tissues were washed and secondary antibodiy from the Vectastain Elite ABC-HRP Rabbit IgG kit was added to the tissue. The tissue was washed and the Vectastain reagent was added. Following the Vectastain reagent, the tissue was subjected to horseradish peroxidase until a noticeable color change was achieved. The tissue was then counterstained with hematoxylin, dehydrated and mounted using Permount. The tissue slides were imaged using a Zeiss microscope at 10X magnification. Alternatively, images across at least three patients per analysis group were captured and analyzed using the Human Protein Atlas [29].

### Cell Proliferation/Viability Assay

Cells were plated at 1e3 cells/well (200μL) in a 96-well plate and allowed to attach overnight. Cells were then treated with conditioned media from stromal cells for 72 hours. For experiments with drug treatment, drugs were added 24 hours after CM treatment. At treatment endpoint, 40μL of the CellTiter 96® AQueous One Solution (Promega) was added to each well. Absorbance readings were measured at 490nm after 3 hours of incubation with reagent using a Spectra Max 190 (Molecular Devices). Absorbance readings are directly proportional to the number of viable cells.

### Incucyte

Cells were trypsinized, pelleted, and resuspended in RPMI medium supplemented with 2% FBS before plating in 200 uL/well into 96-well plates. Mono-cultures received 5e4 cells of cells. Co-cultures received 2.5e4 cells of each respective cell type per well. Two days after seeding, drugs or vehicle controls along with EtBr were prepared at five times the target concentration in 2% FBS-RPMI, and 50uL was added to the designated wells for treatment to obtain the final working concentrations. IncuCyte® Live Cell Analysis Imaging System was used according to manufacturer’s protocol. Four images were taken per well for both the green and red channels. These were collected every 3 hours for the indicated time from time of plating up to 6 days after drug treatment.

### Statistics

All quantified data were plotted and analyzed in GraphPad Prism with ANOVA, Student t test, or nonlinear regression analysis. Data are representative of at least 3 independent experiments and are reported as replicate averages ± SEM, unless otherwise indicated. *, **, *** or **** represent p-values < 0.05, 0.01, 0.001, or 0.0001 respectively, unless otherwise noted.

## RESULTS

### A SNAI2-PEAK1 axis in breast cancer stroma correlates with disease relapse in HER2-positive breast cancer

Since little is known about whether PEAK1 expression or genomic alterations correlate with breast cancer outcomes, we examined the survival of patients across all subtypes in relation to PEAK1 expression levels using the KMPlot database for Kaplan-Meier analysis. This resource enabled assessment of relapse-free survival (RFS), distant metastasis-free survival (DMFS) and overall survival (OS) across more than 3,000 patients [30, 31]. Interestingly, elevated PEAK1 expression across all breast cancer subtypes predicts a significant, though modest, increase in RFS (Figure 1a), while elevated PEAK1 expression in HER2-positive breast cancers showed a striking correlation with decreased RFS suggesting a role for PEAK1 in the relapse of this more aggressive, and often hormone-refractory, breast cancer subtype (Figure 1b). Further, we observed a significant decrease in DMFS across all breast cancer subtypes in patients with high PEAK1 expression (Supplemental Figure 1), consistent with previously reported roles of PEAK1 in metastatic spread of solid tumors [15, 16]. In parallel, we mined data [32] on breast cancer stromal gene expression and discovered that not only was PEAK1 expression significantly higher in malignant breast stroma (Figure 1c), but also that this elevated stromal PEAK1 expression significantly correlated with an increase in disease relapse (Figure 1d).

**Figure 1:**
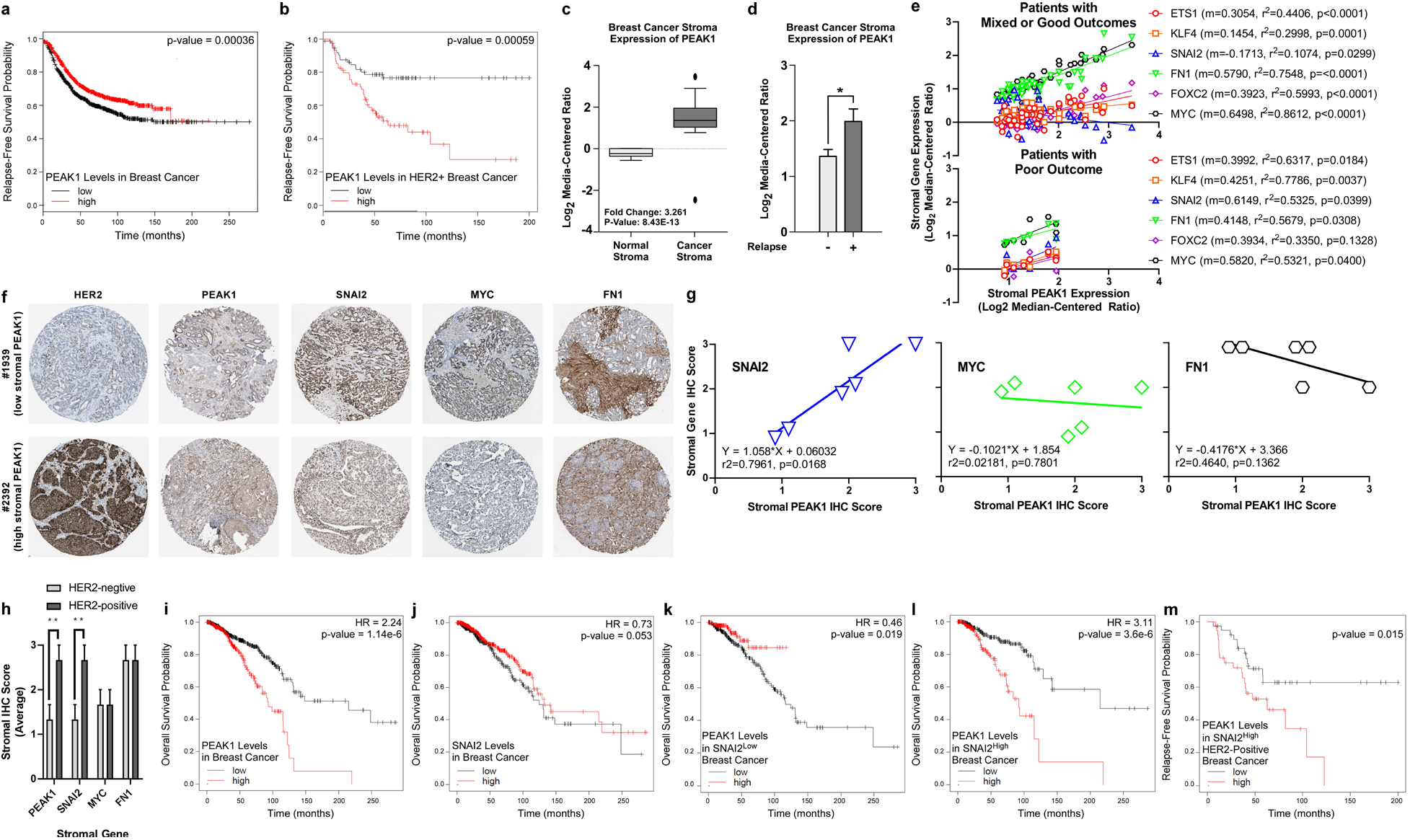
A SNAI2-PEAK1 axis in breast cancer stroma correlates with disease relapse in HER2-positive breast cancer. a. Kaplan-Meier relapse-free survival (RFS) curves for patients with low or high PEAK1 transcript levels across all breast cancer subtypes (n = 1784). b. Kaplan-Meier RFS curves for HER2-positive breast cancer patients with low or high PEAK1 transcript levels (n = 272). c. PEAK1 transcript levels in normal breast stroma and breast cancer stroma across all subtypes (n = 6 and 53, respectively). d. PEAK1 transcript levels in breast cancer stroma of relapse-free patients and those with disease recurrence across all subtypes (n = 42 and 11, respectively). e. Expression correlation and linear regression analyses of PEAK1 and ETS1, KLF4, SNAI2, FN1, FOXC2 and MYC across patients having mixed or good outcomes (top, n = 45) and patients having poor outcomes (bottom, n = 8). f. Representative IHC images for HER2, PEAK1, SNAI2, MYC and FN1 in breast cancer tissue where PEAK1 expression in the stromal compartment is low (top) or high (bottom). g. Stromal IHC score correlation and linear regression analyses of PEAK1 and SNAI2, MYC and FN1 across six breast cancer tissue samples (patient #s 1874, 1939, 2428, 1920, 2392 and 3257). h. Average stromal IHC scores for PEAK1, SNAI2, MYC and FN1 in three HER2-negative patients and three HER2-positive patients. i-j. Kaplan-Meier overall survival (OS) curves for patients with low or high PEAK1 (h) or SNAI2 (i) transcript levels across all breast cancer subtypes (n = 1089). k-l. Kaplan-Meier OS curves for low or high PEAK1 transcript levels in breast cancer patients selected for low (j) or high (k) SNAI2 expression (n = 544). m. Kaplan-Meier RFS curves for low or high PEAK1 transcript levels in HER2-positive breast cancer patient selected for high SNAI2 expression (n = 75). * or ** indicates p-value < 0.05 or 0.01, respectively, as determined by a Student’s T-test.

In order to identify gene networks within breast cancer stroma that may regulate PEAK1 and promote poor survival, we first analyzed how expression patterns for gene signatures corresponding to epithelial (9 genes), mesenchymal (19 genes), stem (4 genes) and mesenchymal stem (15 genes) markers clustered relative to PEAK1 in stromal tissue samples across patient groups previously classified as having poor, mixed or good outcomes [32] (Supplemental Figure 2a). Stromal expression correlation analysis between PEAK1 and six genes that strongly clustered with PEAK1 in the poor outcome group (i.e., ETS1, KLF4, SNAI2, FN1, FOXC2 and MYC) revealed that only the SNAI2-PEAK1 relationship shifted from a significantly negative correlation across patients having mixed or good outcomes to a significantly positive correlation across patients having poor outcomes (Figure 1e). In further support of a SNAI2-PEAK1 stromal cell signaling axis in breast cancer, we noted a significant positive correlation between SNAI2 and PEAK1 protein levels in the stromal compartment of breast cancer samples (Figures 1f-g), and that these SNAI2 and PEAK1 protein levels were specifically expressed at higher levels within the stroma of HER2-positive tumors (Figure 1h) – patterns not observed for either MYC or FN1. In contrast to SNAI2 and MYC, high PEAK1 expression alone does predict poor OS in breast cancer patients (Figures 1i-j and Supplemental Figure 2b). Importantly, however, this prognostic utility of high PEAK1 expression was restricted to patients either expressing high SNAI2 levels (Figures 1k-l) or low MYC levels (Supplemental Figures 2c-d). Finally, we discovered that high PEAK1 expression in HER2-positive breast cancer predicts a significant and further reduction in RFS among patients expressing high levels of SNAI2 transcripts (Figure 1m), further supporting a clinical role for PEAK1 downstream of SNAI2 in the stroma of this aggressive breast cancer subtype.

### SNAI2 and PEAK1 coexpression in breast cancers enriched for mesenchymal stem cell content is prognostically unfavorable

To gain insight into the breast cancer stromal cell subpopulations from within which SNAI2 and PEAK1 may elicit cooperative protumorigenic functions, we mined clinical data for relationships between high coexpression of SNAI2 and PEAK1 and poor overall survival across patient tissues enriched for innate immune, adaptive immune or mesenchymal stem cell content. As shown in Figures 2a-b, high expression of both SNAI2 and PEAK1 did not predict overall survival probability among breast cancer patients reporting enrichments in either innate or adaptive immune cell content. Notably, high expression of both SNAI2 and PEAK1 predicted significantly lower overall survival among patients with mesenchymal stem cell (MSC) content (Figure 2c). In contrast, high PEAK1 expression levels among patients with high SNAI2 expression did not offer any prognostic significance in patients with decreased MSC content (Figure 2d), supporting a cooperative protumorigenic role for SNAI2 and PEAK1 among mesenchymal stromal cell types in the breast cancer microenvironment.

**Figure 2:**
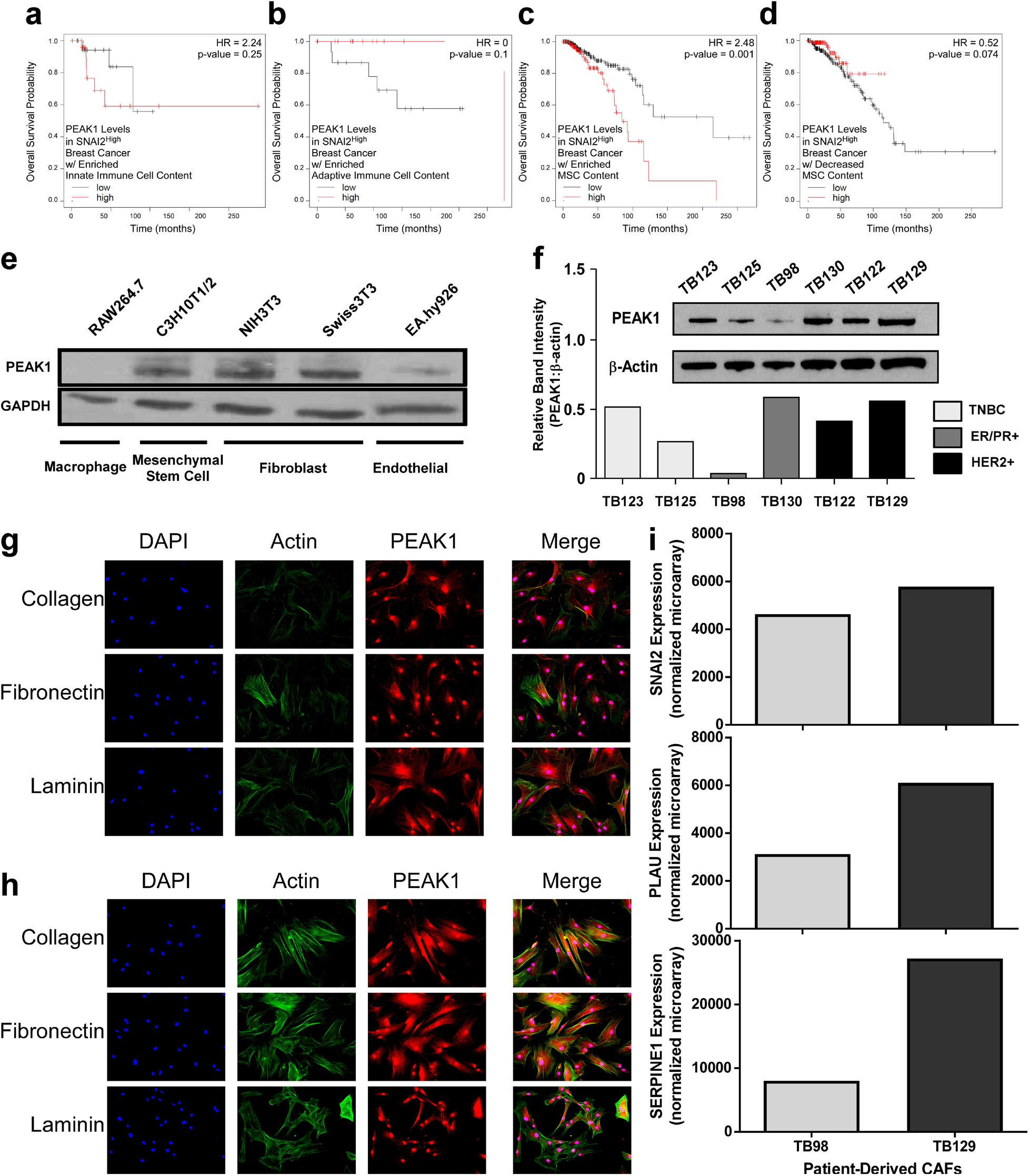
Co-expression of SNAI2 and PEAK1 in breast cancers enriched for mesenchymal stem cell content is prognostically unfavorable. a-c. Kaplan-Meier OS curves for low or high PEAK1 transcript levels in breast cancer patients selected for high SNAI2 expression and enriched innate immune (a), adaptive immune (b) or mesenchymal stem cell contents (n = 45, 19 and 382, respectively). D Kaplan-Meier OS curves for low or high PEAK1 transcript levels in breast cancer patients selected for low SNAI2 expression and enriched mesenchymal stem cell content (n = 381). e. Western blot for PEAK1 and GAPDH in total lysates from the indicated non-tumorigenic cell lines. f. Western blot and relative band intensity for PEAK1 and β-actin in total lysates from the indicated patient-derived breast cancer associated fibroblasts. g-h. immunofluorescence for nucleus (DAPI), filamentous actin (Phalloidin) and PEAK1 in TB98 (g) and TB129 (h) CAF lines plated onto 5 ug/mL collagen, fibronectin or laminin substrates. i. Normalized expression of SNAI2, PLAU and SERPINE1 in the indicated TB CAF lines obtained from the GEO database using the GSE37614 dataset.

By evaluating PEAK1 expression across non-tumor cell lines that are naïve to the breast cancer microenvironment but which share similarities to those found within this stromal context (i.e., 3 fibroblast-like, 1 endothelial and 1 innate immune cell lines), we further established that PEAK1 expression was primarily detected within fibroblast-like cell types (Figure 2e). These data were further supported by analyzing the expression of PEAK1 across a subset of patient-derived cancer associated fibroblasts (CAFs) (i.e., two from each breast cancer subtype) previously isolated and transcriptomically profiled [27]. While CAFs from ER-positive displayed the most heterogeneity for PEAK1 expression levels, CAFs from all breast cancer subtypes expressed PEAK1 (Figure 2f). At the subcellular level, PEAK1 localized with the actin cytoskeleton in both the TB98 (from ER+ breast cancer) and TB129 (from HER2-positive breast cancer) CAF lines independent of extracellular matrix (ECM) substrate (Figure 2g-h). Finally, analysis of SNAI2 and two other mesenchymal stromal cell genes (i.e. PLAU and SERPINE1) revealed that their collective expression to be higher in the TB129 CAF line relative to the TB98 line (Figure 2i) – data consistent with PEAK1 expression patterns.

### Xenografting patient-derived CAFs or MSCs with HER2-positive breast cancer cells increases primary tumor mass

We next asked whether PEAK1-expressing CAFs or MSCs could affect breast tumor growth and progression *in vivo*. Using the chicken chorioallantoic membrane (CAM) xenograft model [28, 33, 34], we implanted either HER2-positive BT474 or ER-positive MCF7 breast cancer cells onto the CAM of 10 day old chicken embryos and allowed them to grow for an additional 7 days before harvesting the primary CAM tumor and isolating genomic DNA from lung and brain tissues to quantify alu sequences as a measure of systemic metastasis (Figure 3a). As shown in Figures 3b-c, BT474 cells were able to make primary CAM tumors of greater mass when xenografted together with either the TB122 CAFs or C3H10T1/2 MSCs, although neither the CAF- nor MSC-containing xenografts displayed a measurable difference in dissemination to lung or brain tissues (Figure 3d). Similarly, MCF7 cells xenografted alone, together with TB130 CAFs or together with TB130 CAFs after *in vitro* pre-incubation with TB130 CAF conditioned media (CM) were able to form primary tumors and the lung and brain tissue from these animals did not display any macroscopic metastatic nodules (Supplemental Figure 3a). Quantification of MCF7 CAM tumor mass from these animals demonstrated that the TB130 CAFs were able to potentiate MCF7 tumor growth *in vivo* (Supplemental Figure 3b). However, these CAF-containing xenografts did not affect the ability of MCF7 cells to metastasize to either the lung or brain (Supplemental Figure 3c).These results are consistent with previous findings demonstrating a supportive role for MSCs on solid tumor growth *in vivo* [35], and establish this system as a means to interrogate the role of PEAK1 in MSC-mediated HER2-positive breast cancer progression and targeted therapy response.

**Figure 3:**
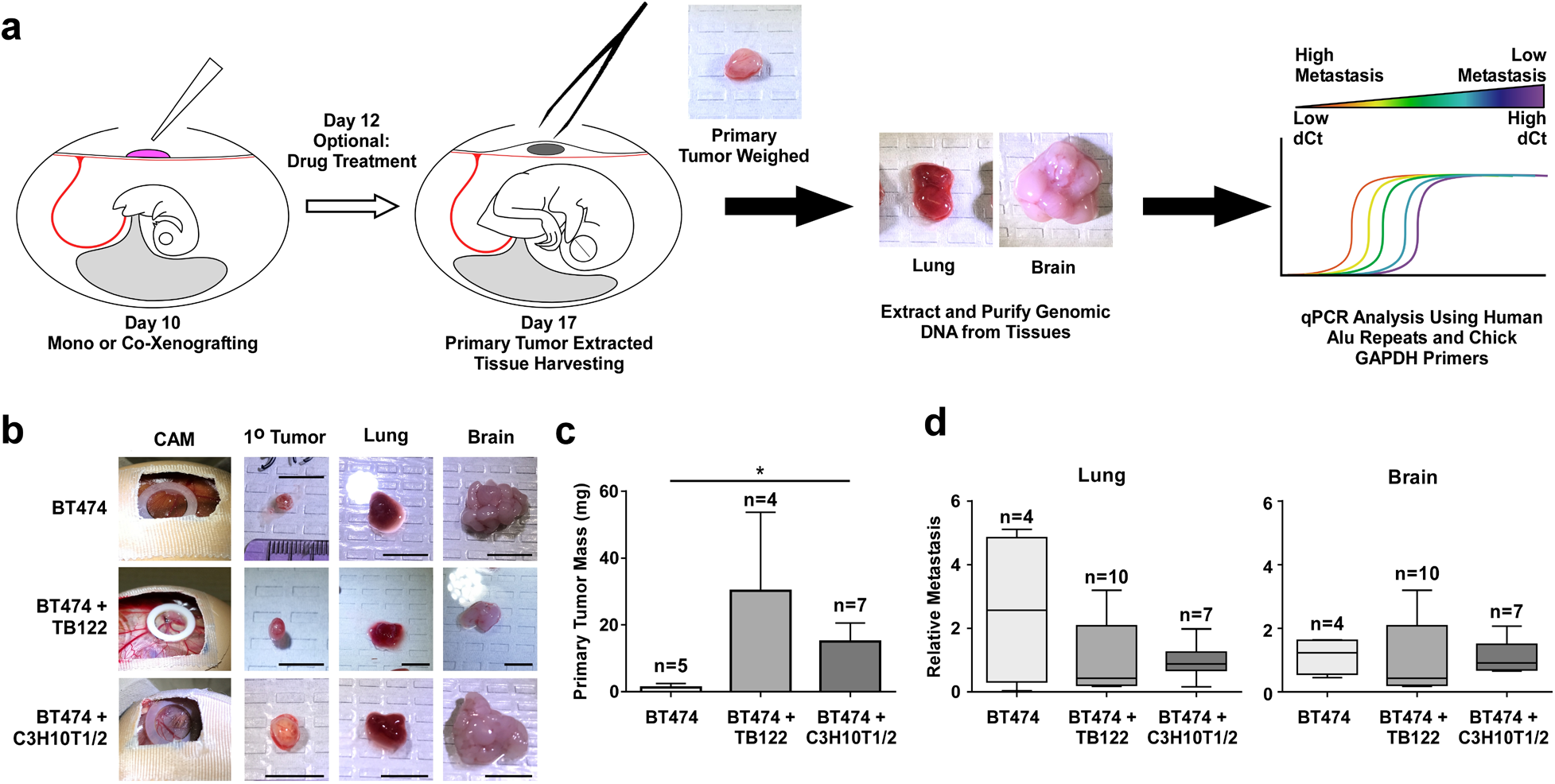
Xenografting patient-derived CAFs or MSCs with HER2-positive breast cancer cells increases primary tumor mass. a. Schematic of the chorioallantoic membrane (CAM) xenograft system using chicken (Gallus gallus) embryos together with human tumor cells and end point analysis of whole tissue genomic DNA by qPCR for human alu repeats and host chicken Gapdh levels. The method has been modified from the original assay system (Zjlistra et al. 2002 Cancer Research) to enable a 5-day drug treatment regiment beginning at 2 days post-xenograft. b. Indicated images of BT474 cells, BT474 cells + TB122 CAFs or BT474 cells + C3H10T1/2 mesenchymal stem cells. Scale bar = 1 cm. c. Quantified primary tumor mass of experiment in (b). d. Relative metastasis of BT474 cells in the lung (left) and brain (right) of experiment in (b). * indicates a p-value < 0.05 as determined by a One-Way ANOVA w/ multiple comparisons test.

### Knockdown of PEAK1 in MSCs abrogates their ability to promote tumorigenesis, intratumoral αSMA expression, lapatinib resistance and lapatinib-induced brain metastasis

To assess the function of PEAK1 in mediating these intercellular protumorigenic functions of MSCs, we generated a panel of stable shRNA C3H10T1/2 MSC derivatives containing either a scramble control shRNA construct (shScr) or one of five PEAK1-targeting shRNAs (shP1) (Figure 4a). As before with the parental C3H10T1/2 MSCs, xenografting the shScr MSCs with the BT474 cells significantly increased primary tumor mass – an effect that was abrogated by PEAK1 knockdown using two unique shRNA constructs (Figure 4b). Notably, the shScr MSC containing BT474 xenograft tumors showed regions enriched for alpha smooth muscle actin (αSMA) staining and vascular-like structures (Figure 4c).

**Figure 4:**
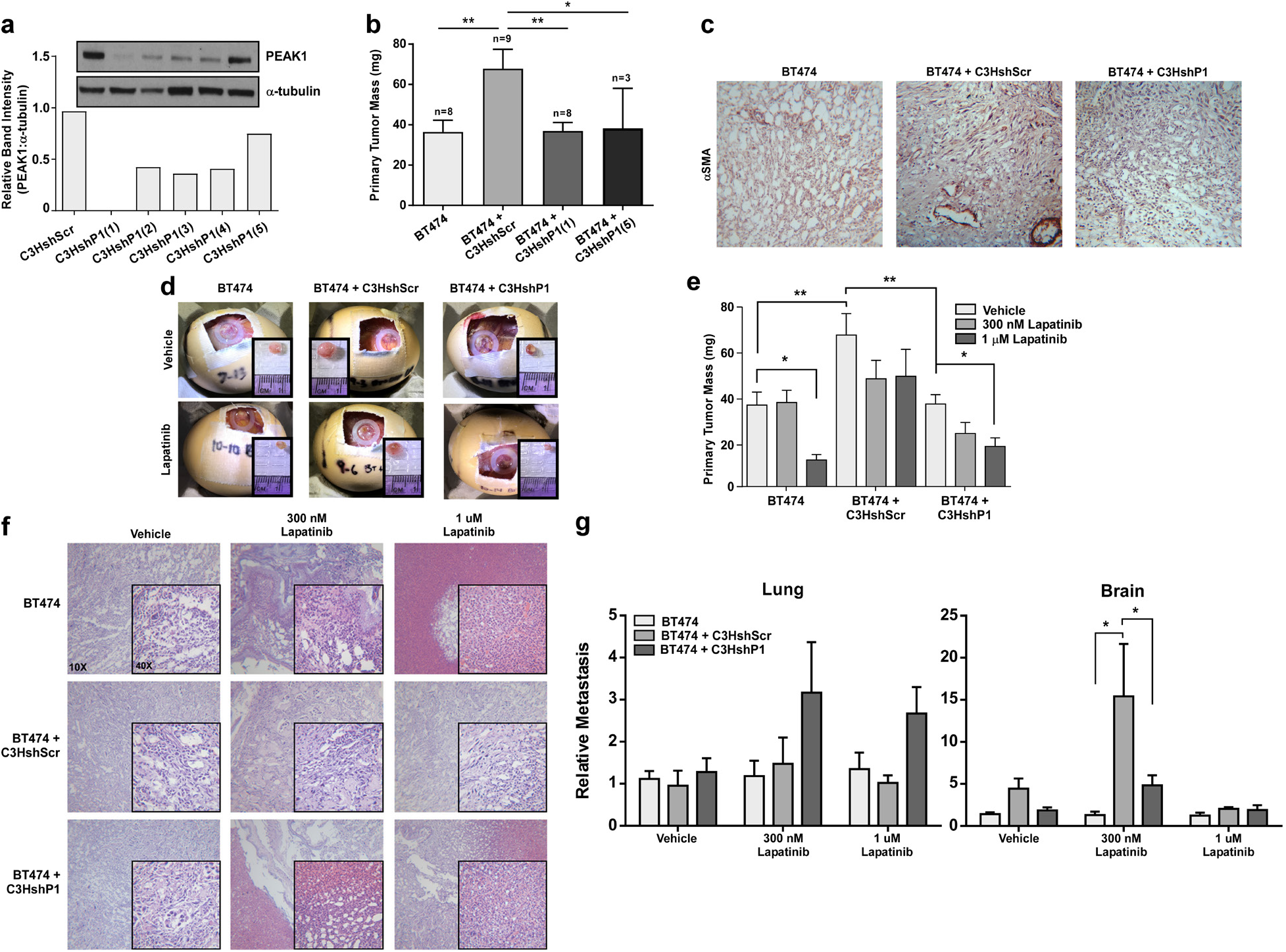
Knockdown of PEAK1 in MSCs abrogates their ability to promote tumorigenesis, intratumoral αSMA expression, lapatinib resistance and lapatinib-induced brain metastasis. a. Western blot and relative band intensity quantification for PEAK1 and a-tubulin levels in shScramble control (shScr) and 5 different PEAK1 targeting (shP1) shRNA derivatives of C3H10T1/2 mesenchymal stem cells. Unless otherwise noted, shP1(1) construct is used throughout experiments. b. Quantified primary tumor mass of CAM xenograft assay using BT474 cells only or BT474 cells xenografted together with the C3HshScr, C3HshP1(1) or C3HshP1(5) cells. c. a-smooth muscle actin staining of stromal tissue in CAM tumors from (b). d. Assay endpoint images for CAM xenograft experiment using the same cell combinations as in (b) with either vehicle control or 1 uM lapatinib treatments. e. Quantified primary tumor mass of experiment in (d) including the treatment condition of 300 nM lapatinib. f. Hematoxylin and eosin staining of CAM tumor tissue in (d). g. Relative metastasis of BT474 cells in the lung (left) and brain (right) of experiment in (d). * or ** indicates a p-value < 0.05 or 0.01, respectively, as determined by a One-Way or Two-Way ANOVA w/ multiple comparisons post-test.

Given that previous work has described a role for the spatial proximity of αSMA-positive stromal fibroblasts in promoting lapatinib resistance and proliferation of residual HER2-positive breast cancer cells [4], we analyzed whether these MSCs could render BT474 tumors resistant to lapatinib treatment *in vivo* and whether any observed effects might require PEAK1 expression. Notably, shScr MSCs rendered BT474 cells less sensitive to lapatinib *in vivo*. Furthermore, BT474 xenografts containing MSCs with the PEAK1-targeting shRNAs responded to lapatinib as though there were no MSCs xenografted onto the CAMs with the breast cancer cells (Figures 4d and 4e). In agreement with these data, hematoxylin and eosin (H&E) stained primary CAM tumors revealed a high degree of dead tissue around the periphery of tumors generated from either BT474 cells alone or BT474 xenografts containing MSCs with the PEAK1-targeting shRNAs at both lapatinib doses (Figure 4f). In contrast, tumors from BT474 cells xenografted with the shScr MSCs lacked any necrotic regions within the tumors even at the highest lapatinib dose (Figure 4f). Interestingly, while PEAK1-expressing MSCs were unable to induce metastasis alone, the presence of these cells in the primary tumors treated with intermediate doses of lapatinib enabled the BT474 cells to escape and metastasize to the brain at a 15-fold greater frequency when compared to xenografts of the BT474 cells alone or BT474 cells and shP1 MSCs (Figure 4g).

### MSC expression of PEAK1 protects neighboring breast cancer cells from lapatinib-induced cytotoxicity

To elucidate potential mechanisms by which these stromal cells elicit their tumor- and lapatinib resistance-promoting functions, we established a co-culture system to further evaluate whether MSCs or breast fibroblasts could promote breast cancer cell expansion and resistance to lapatinib *in vitro* (Figure 5a). Co-seeding either MSCs or AR22 breast fibroblasts together with H2B-eGFP labeled BT474 cells at high density and a 1:1 ratio established monolayer co-cultures in which the breast cancer cells formed islands surrounded by fibroblasts (Figure 5a and Supplemental Figure 4a) – a two dimensional architecture similar to that observed in patient tumors. We then used this system in combination with IncuCyte imaging to evaluate both the number of eGFP-positive and EtBr-positive (membrane-compromised and dying) breast cancer cells during time-course lapatinib dose-response experiments (Figure 5a). As shown in Supplemental Figure 4b, mono-cultures of either the MSCs or fibroblasts did not respond to increasing lapatinib doses as measured by EtBr uptake while the BT474 mono-cultures did, demonstrating that this targeted therapy displays specific cytotoxicity to HER2-overexpressing breast cancer cells. Interestingly, while co-culture of BT474 cells together with MSCs was able to both increase the basal number of BT474 cells (Figure 5b) and reduce lapatinib cytotoxicity (Figure 5c), the breast fibroblasts seemed to selectively reduce lapatinib cytotoxicity (Figure 5c and Supplemental Figure 4b). Using the shScr and shP1 MSC derivatives in this co-culture assay revealed that PEAK1 expression specifically mediates the ability for MSCs to protect neighboring breast cancer cells against lapatinib-induced cytotoxicity (Figure 5d).

**Figure 5:**
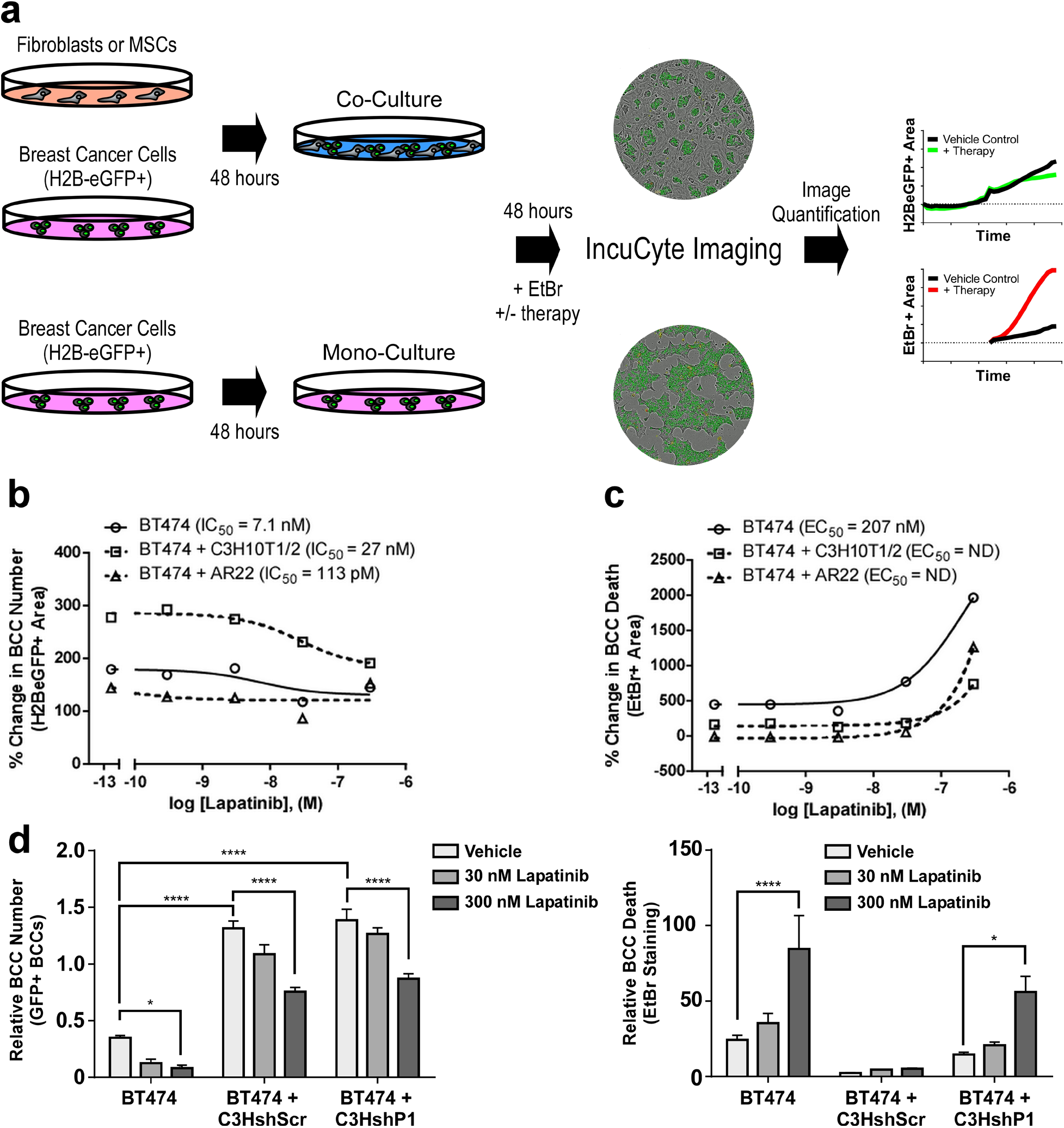
MSC expression of PEAK1 protects neighboring breast cancer cells from lapatinib-induced cytotoxicity. a. Schematic of breast cancer cell mono- or breast cancer cell-CAF/MSC co-culture using non-labeled stromal fibroblasts and H2B-eGFP+ BT474 breast cancer cells for downstream analysis of breast cancer cell number and death using the IncuCyte imaging system over 96 hours (48 hours pre-incubation and 48 hours incubation with therapy and EtBr). b and c. Endpoint dose-response curves for lapatinib effects on breast cancer cell number (b) or cell death (c) in the indicated breast cancer cell and stromal fibroblast culture combinations. d. Quantification of tumor cell number (left) and EtBr uptake (right) at assay endpoint for BT474 cells alone or co-cultured with the shScr or shP1(5) derivatives of C3H10T1/2 cells and treated with vehicle control or the indicated dose of lapatinib. * of **** indicates a p-valule of 0.05 or 0.0001, respectively as determined by a Two-Way ANOVA with multiple comparisons post-test.

### PEAK1 expression in MSCs drives the production of secreted factors that promote breast cancer cell proliferation/survival and lapatinib resistance in vitro

To determine whether MSCs and CAFs can promote breast cancer expansion and/or targeted therapy resistance cell via paracrine mechanisms, we tested conditioned media (CM) from these cell types for its ability to promote breast cancer cell expansion and/or lapatinib resistance *in vitro* (Figure 6a). CM collected from CAFs derived from either HER2-positive (Figure 6b) or ER-positive (Supplemental Figure 5a) breast cancers were able to potentiate BT474 or MCF7 cell growth, respectively. Notably, by using shRNA MSC derivatives, we report that PEAK1 is necessary to produce secreted factors into MSC CM that potentiate BT474 (Figure 6c) or mouse Py230 (Supplemental Figure 5b) cell growth. Finally, we report that MSC expression of PEAK1 was necessary for MSC-derived CM to promote BT474 cell resistance to lapatinib (Figure 6d). Taken together, these data demonstrate that PEAK1 expression is required for MSCs and CAFs to produce and/or secrete soluble factors that can promote proliferation/survival and lapatinib resistance in breast cancer cells *in vitro*.

**Figure 6:**
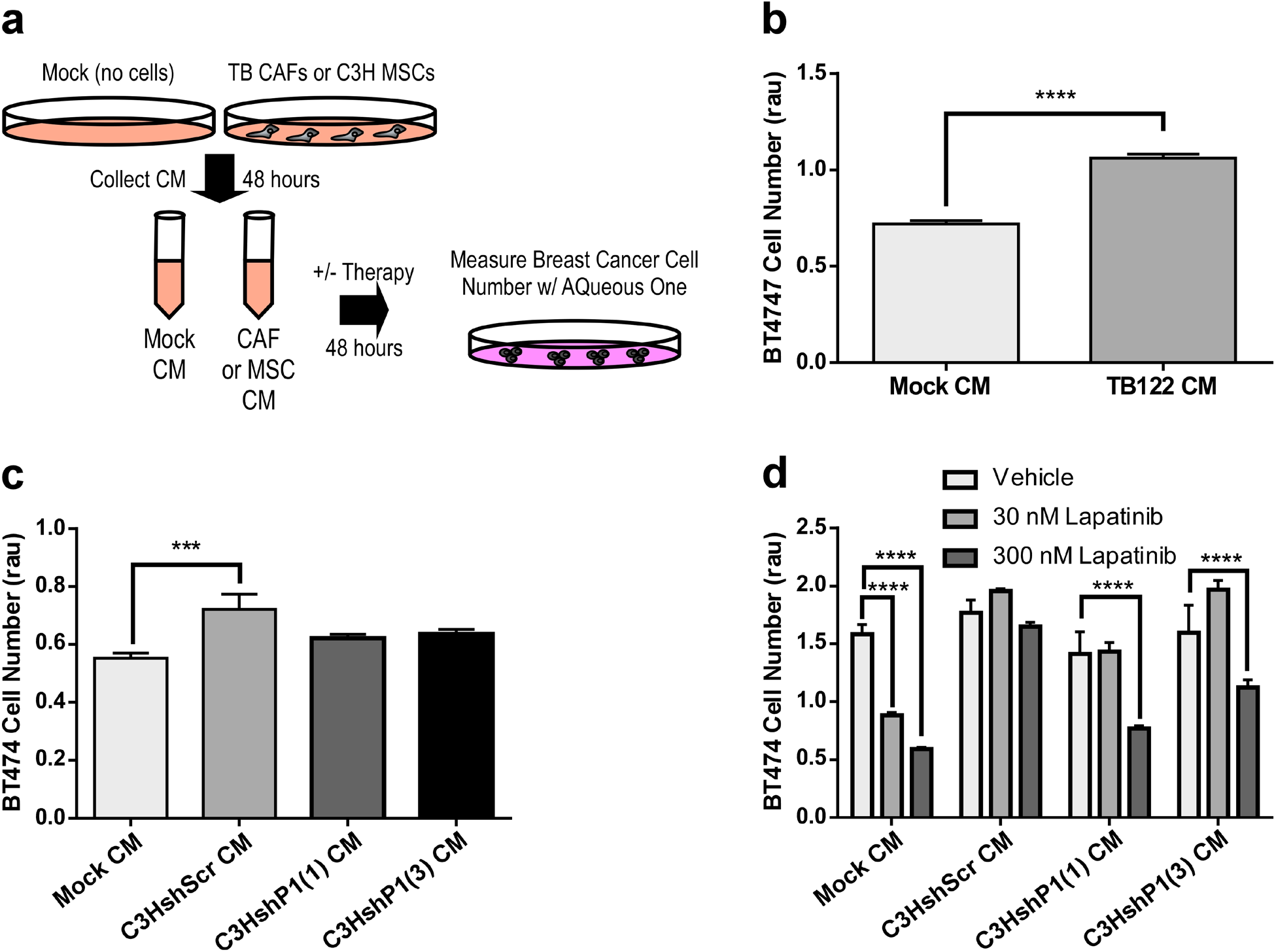
PEAK1 expression in MSCs drives the production of secreted factors that promote breast cancer cell proliferation/survival and lapatinib resistance *in vitro*. a. Schematic for generating TB CAF or C3H MSC conditioned media (CM) for analysis on breast cancer cell growth/survival over 48 hours in vitro. b. Cell viability analysis of BT474 cells treated with mock or TB122 CM. c. Cell viability analysis of BT474 cells treated with mock CM or CM from the indicated shRNA derivatives of C3H10T1/2 cells. d. Cell viability analysis of BT474 cells treated with mock CM or CM from the indicated shRNA derivatives of C3H10T1/2 cells and treated with vehicle control or the indicated dose of lapatinib. *, ***, or **** indicates a p-value < 0.05, 0.001 or 0.0001, respectively, as determined by a One-Way or Two-Way ANOVA w/ multiple comparisons post-test.

### A PEAK1-dependent cytokine profile in MSCs supports PI3K/Akt signaling and lapatinib resistance in HER2-positive breast cancer cells

To identify the factors that PEAK1 regulates within MSCs, we performed semi-quantitative protein array analysis targeting 308 protein antigens in lysates from the shScr and two unique shP1 MSC derivatives (Figure 7a). PEAK1 knockdown led to a greater than 2-fold decrease in 5 proteins (GDF5, CCR4, INHBA, GRH and CCL4) and a greater than 1.8-fold increase in 7 proteins (PDGFRA, CSF1, HGFR, Frizzled-6, VEGFA, PF4 and TGFB3) (Dataset 1). As shown in Figure 7b, six PEAK1-dependent soluble factors (four of which are members of the transforming growth factor beta, TGFβ, Superfamily) met the 95% confidence interval cut-off criteria for further analysis. Using the Core Expression Analysis function within the Ingenuity Pathway Analysis (IPA) platform, interactome networks were generated for PEAK1 and PEAK1-suppressed proteins (i.e., TGFB3, VEGFA and CSF1) (Figure 7c, left) or PEAK1-induced proteins (i.e., CCL4, INHBA and GDF5) (Figure 7c, right). Interestingly, while the PEAK1-suppressed genes/proteins generated an interactome that converged on ERK1/2-regulating genes/proteins (Figure 7c, left), the PEAK1-induced genes/proteins generated an interactome which converged on AKT1-regulatinng genes/proteins (Figure 7c, right). Further analysis of these interactome nodes for enriched canonical signaling pathways (Figure 7d), diseases and disorders (Figure 7e), molecular and cellular functions (Figure 7f), and system development and function (Figure 7g) revealed that the PEAK1-induced interactome focuses on ErbB signaling, cancer and cell-to-cell signaling and interaction (Figures 7d-f). To determine if Akt signaling in HER2-positive breast cancer cells may be an important mediator of PEAK1-dependent MSC-induced lapatinib resistance, we assessed Akt phosphorylation levels in co-cultures of shRNA MSC derivatives and the BT474 cells after a 24 hour treatment of vehicle control or an intermediate dose of lapatinib. While Akt phosphorylation was predominantly restricted to H2BeGFP-positive BT474 cells and lapatinib nearly abolished Akt phosphorylation in the mono-culture condition, co-cultures of BT474 cells and shScr MSCs increased basal Akt phosphorylation in the BT474 cells and sustained it following lapatinib treatment – effects for which PEAK1 expression in the MSCs was required (Figure 7h-i).

**Figure 7:**
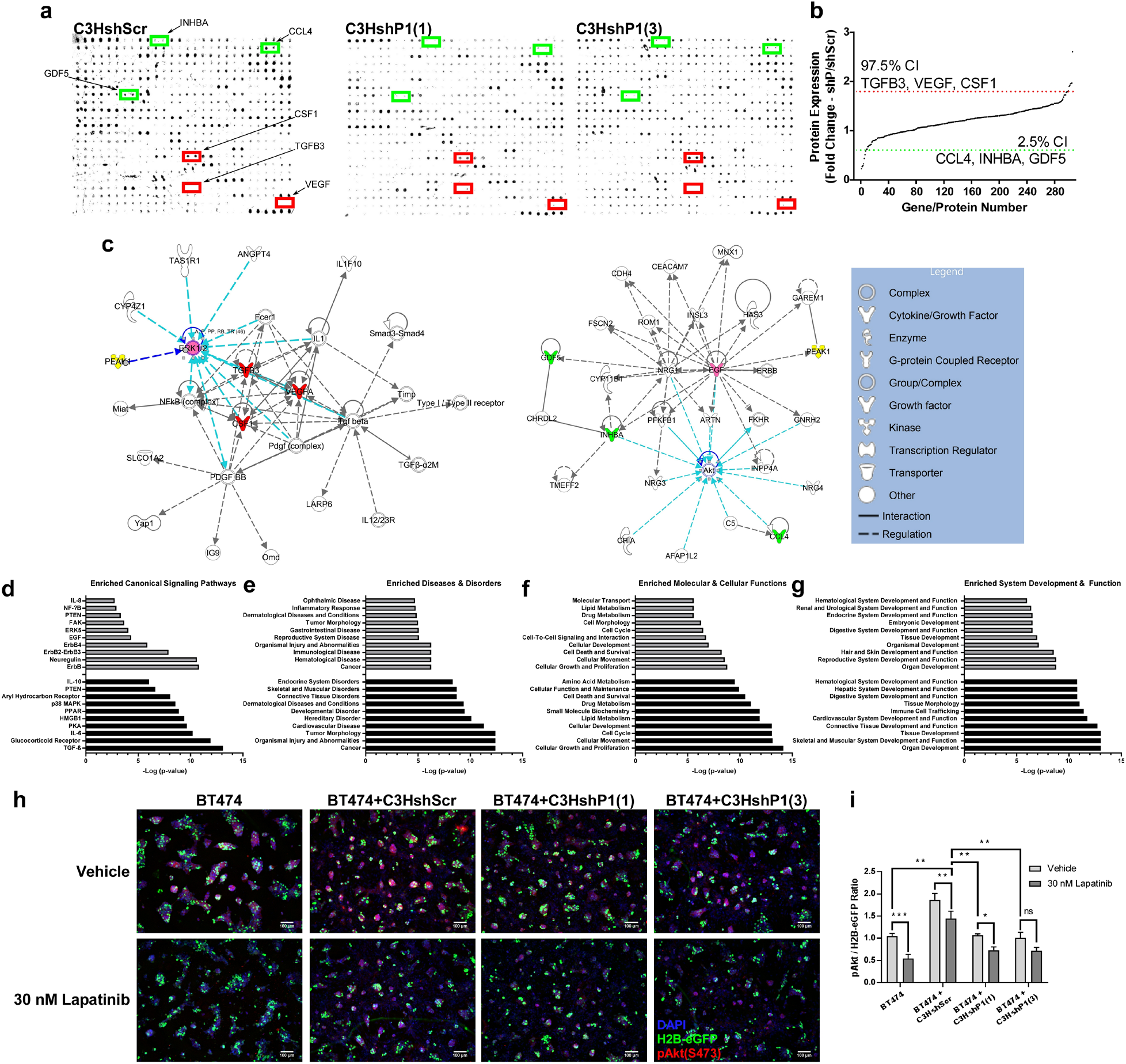
A PEAK1-dependent cytokine profile in MSCs supports PI3K/Akt signaling and lapatinib resistance in HER2-positive breast cancer cells. a. Representative slide scan images from the semi-quantitative mouse antibody array 308 (L-308) following incubation and reactivity with total cell lysates prepared from the shScramble and indicated PEAK1-specific shRNA derivatives of the C3H10T1/2 MSCs. b. Quantification of ranked protein expression across the 308 array antigens with confidence intervals set to identify antigen expression changes up (red line) and down (green) with a p-value < 0.05. c Ingenuity Pathway Analysis (IPA)-generated networks for PEAK1, TGFB3, VEGFA and CSF1 (left) and PEAK1, CCL4, INHBA and GDF5 (right). Yellow = PEAK1, Red = PEAK1-suppressed genes/proteins, Green = PEAK1-induced genes/proteins and Blue lines = molecular convergence. d-g. Enriched canonical signaling pathways, diseases & disorders, molecular & cellular functions and system development & function for the IPA networks in c-left (PEAK1-suppressed, bottom/black) and c-right (PEAK1-induced, top/grey). h. Representative widefield immunofluorescence for GFP (H2B-eGFP-positive BT474 cells, green), DAPI (nucles, blue) and phospho-Akt (red) levels in the mono- or co-culture BT474-C3H10T1/2 cell system outlined in Figure 5a. Indicated samples were treated with lapatinib for 24 hours prior to fixing/staining/imaging.

### INHBA/Activin A expression is upregulated in malignant breast stroma, correlates with PEAK1 levels in breast cancer tissue and predicts poor survival in HER2-positive patients or PEAK1-high breast cancers with enriched MSC content

We next sought to determine whether there was any clinical relevance of these PEAK1-regulated MSC gene/proteins (i.e., TGFB3, VEGFA, CSF1, CCL4, INHBA and GDF5) in relation to their concurrent upregulation with PEAK1 in HER2-positive breast cancers enriched for MSC content. As with PEAK1 (Figure 1), we analyzed expression profiles for these genes across two independent studies investigating normal breast stroma and malignant breast stroma [5, 32] (data retrieved from Oncomine). Notably, the transcripts of PEAK1-suppressed MSC factors (i.e., TGFB3, VEGFA and CSF1) were significantly lower within breast cancer stroma (Figure 8a, left three graphs). In contrast, the factors whose expression was positively regulated (i.e., induced) by PEAK1 in MSCs (i.e., CCL4, INHBA and GDF5) displayed significantly higher transcript levels in breast cancer stroma across both studies, with INHBA (ActivinA) showing the greatest average fold-change increase in malignant over normal breast stroma (Figure 8a, right three graphs). Further analysis of the mRNA expression relationship between PEAK1 and each of these six genes in breast cancer patient tissues using Cancer BioPortal [36] revealed that PEAK1 and INHBA transcripts showed the most significant positive correlation (Figure 8b). We next assessed the relationship between expression levels for each of the six PEAK1-regulated genes and RFS survival across all breast cancer subtypes (Figure 8c) and HER2-positive breast cancer (Figure 8d). While INHBA levels had no prognostic significance across breast cancers from all subtypes, increased INHBA expression correlated strongly with decreased RFS within HER2-positive breast cancer (Figure 8d), as did TGFB3 and VEGFA. Finally, analysis of how PEAK1 expression affects OS in breast cancer patients with high expression for each of the six PEAK1-regulated genes together with enriched MSC content, identified co-upregulaltion of INHBA/ActivinA and PEAK1 as the most significant predictor of poor outcome (hazard ratio = 3.27 and median OS = 75 months) (Figure 8e). Taken together, these data support a role for PEAK1-mediated INHBA/ActivinA expression or proteostasis within the mesenchymal stromal compartment as a mechanism that promotes progression and poor outcomes in HER2-positive breast cancer.

**Figure 8:**
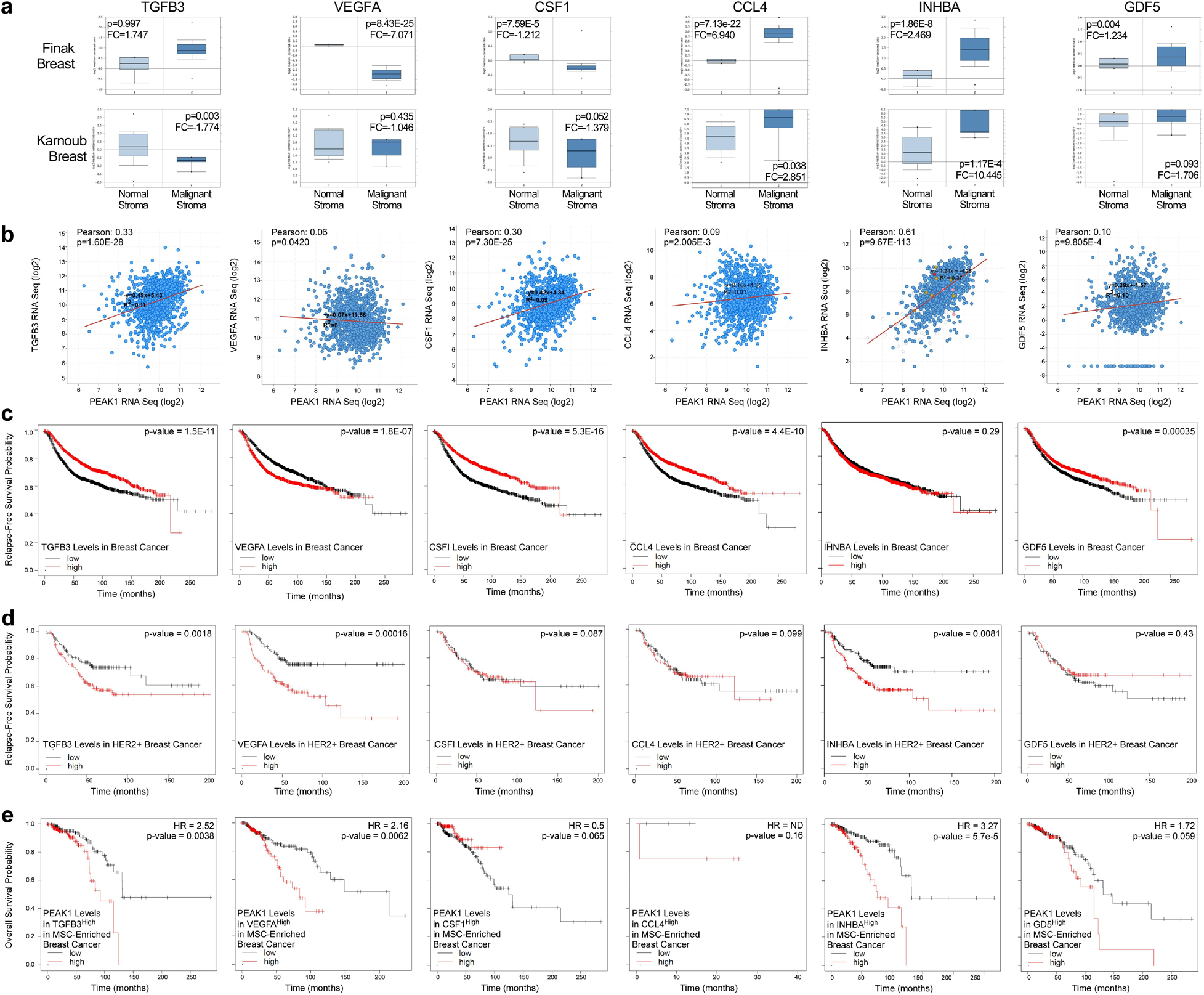
INHBA/Activin A expression is upregulated in malignant breast stroma, correlates with PEAK1 levels in breast cancer tissue and predicts poor survival in HER2-positive patients or PEAK1-high breast cancers with enriched MSC content. a. Relative mRNA expression for the indicated PEAK1-dependent cytokines in normal versus malignant stroma as reported in the indicated studies. b. Expression relationship for TGFB3, VEGFA, CSF1, CCL4, INHBA and GDF5 versus PEAK1 mRNA levels in breast cancer patients. c. Kaplan-Meier Relapse-Free Survival (RFS) curves of breast cancer patients across all subtypes in relation to low or high expression of the indicated cytokines (n = 3951). d. Kaplan-Meier RFS curves of breast cancer patients with HER2-positive disease in relation to low or high expression of the indicated cytokines (n = 252). e. Kaplan-Meier Overall Survival (OS) curves for high and low PEAK1 in patients enriched for MSC content and selected for high expression of the indicated cytokine (n = 380, 392, 382, 12, 350 and 382, respectively).

## DISCUSSION

Here, we identify a pharmacologically targetable PEAK1-dependent and SNAI2-associated stromal cell non-autonomous mechanism through which neighboring HER2-positive breast cancer cells increase in PI3K/Akt signaling activity, acquire lapatinib resistance and metastasize to the brain (Supplemental Figure 6). It is well-documented that targeted therapy resistance of HER2-positive breast cancer strongly associates with the onset of brain metastasis [37]. Importantly, this is the first demonstration that PEAK1 promotes malignant phenotypes from within cancer associated non-tumor cell types. Consistent with our findings that PEAK1 expression predicts poor patient prognosis in the context of high SNAI2 expressing breast cancers (Figure 1l), recent work has reported that reprogramming of stromal fibroblasts by SNAI2 contributes to solid tumor progression and that this occurs concurrent with upregulated stromal PEAK1 transcript levels [38].

Marusyk and colleagues demonstrated that co-culturing or xenografting stromal fibroblasts together with HER2-positive breast cancer cells sustains Akt phosphorylation in the presence of lapatinib treatment [4]. A cancer cell intrinsic role for Akt signaling in promoting resistance of HER2-positive breast cancer to lapatinib treatment was earlier reported from studies of HER2-positive breast cancer xenografts in which engineered tumor cells overexpressed PIK3CA [39]. Resulting tumor tissues had increased Akt phosphorylation levels and developed lapatinib resistance – phenotypes that could be overcome by PI3K inhibition [39]. In this regard, we demonstrate that high PEAK1 expression in HER2-positive breast cancer patient tissues predicts increased disease relapse (Figure 1b) and that this association is specific to HER2-positive breast cancer tissues high in SNAI2 expression (Figure 1m) and mesenchymal stem cell (MSC) content (Figure 2c). Notably, we go on to demonstrate that MSC expression of PEAK1 is necessary for their ability to promote lapatinib resistance, HER2-positive breast cell expansion *in vitro* and *in vivo* (Figures 4-6) and sustained Akt phosphorylation within neighboring breast cancer cells following lapatinib treatment (Figures 7h-i).

A role for MSCs in the breast cancer microenvironment as effectors of tumor growth and metastasis has been previously established [5, 40, 41]. It is interesting that our *in vivo* results did not demonstrate that basal PEAK1-dependent tumor-promoting MSC functions could increase spontaneous metastasis of either HER2-positive BT474 (Figure 3d) or ER-positive MCF7 (Supplemental Figure 3c) cells to lung or brain tissues. However, we note that previous work by Karnoub and colleagues did not evaluate the effect of MSCs on metastasis of HER2-positive breast cancer cells [5] and that the MCF7 cells used in these initial spontaneous metastasis studies were Ras-overexpressing, possibly rendering them more aggressive and susceptible to exogenous factors such as CCL5 that was reported to be induced in these MSCs following their reprogramming by breast cancer cells [5]. Notably, we observed that MSC expression of PEAK1 was required for MSCs to induce metastatic spread of HER2-positive breast cancer cells to the brain in animals treated with lapatinib (Figure 4g). We also noted that the primary tumors in these animals showed no signs of cell death, even at the highest doses of lapatinib (Figure 4f). While the specific cellular processes and/or molecular machinery governing these effects will require further characterization, one possibility is that stromal expression of PEAK1 enhances tumor vascularization. This is consistent with both our observation that HER2-positive breast cancer cells xenografted with MSCs displayed increased expression of alpha smooth muscle actin (αSMA) staining and vascular architecture in a PEAK1-dependent manner (Figure 4c) and recent work reporting a role for PEAK1 during angiogenesis [42].

Previous analyses of cell line xenografts in mice and patient tumor tissue revealed that lapatinib treatment leads to a decreased distance between αSMA-positive stromal fibroblasts and proliferating HER2-positive breast cancer cells [4], implicating juxtacrine and/or distance-dependent paracrine signaling mechanisms such as those used by morphogens. Interestingly, we observed that PEAK1-dependent MSC-induced protection of HER2-positive breast cancer cells against lapatinib could occur in both a co-culture system and by treating breast cancer cells with conditioned media from naïve MSCs (i.e., MSCs not previously exposed to breast cancer cells) (Figures 5-6). The possibility that these MSC-driven cytoprotective effects require one or more secreted factors is supported by our identification of six secreted/soluble proteins (i.e., TGFB3, VEGFA, CSF1, CCL4, INHBA and GDF5) that were expressed by MSCs in a PEAK1-dependent manner (Figures 7a-b). However, it remains to be determined whether PEAK1 regulates the transcription or translation/proteostasis of these genes/proteins and whether PEAK1 dimerization or actin/focal adhesion associations may influence the expression of these factors [43]. Nonetheless, it is enticing to note that TGFBs/Inhibins/Activins have well-established morphogen roles during normal development [44], and that previous work has reported increased INHBA/ActivinA activity at the leading edge of HER2-positive breast tumors [45] and that follistatin (an AcitivinA antagonist) can suppress HER2-positive breast cancer metastasis [46]. These results together with our findings that PEAK1 expression predicts low median overall survival in breast cancer patients with high INHBA expression and enriched for MSC content (Figure 8e), further support a role for PEAK1-dependent INHBA expression as a mechanism by which stromal MSCs support HER2-positive breast cancer progression and therapy resistance.

While it will be important for future studies to identify which combination of PEAK1-dependent factors expressed from within the HER2-positive breast cancer microenvironment are necessary and/or sufficient to induce tumorigenesis, the studies here establish PEAK1 as an critical tumor stromal regulatory node that works in concert with SNAI2 an INHBA to promote therapy resistance, metastasis and poor patient outcomes.

## ACKNOWLEDGEMENTS

We thank Dr. Joan Brugge for sponsorship of Dr. Jonathan Kelber’s ASCB-MAC Visiting Professorship and helpful discussions during the experiment design and manuscript writing/revising phases. We thank Dr. Lesley Ellies for providing Py230 cells for experiments in this work. We also thank Mr. Cameron Geller of the Developmental Oncogene Laboratory at California State University Northridge for his valuable feedback during the writing/revising of this manuscript. This work was supported by California State University Northridge College of Science and Mathematics; Wellcome Trust Centre for Cell-Matrix Research at the University of Manchester; Ludwig Center at Harvard Medical School; Sidney Stern Memorial Trust; Sutter family; NIH NCI grant R00CA222554 (to I.K.Z.); DOD grant W81XWH-14-1-0222 (to I.K.Z); Cancer Research UK programme grant C13329/A21671 (to M.J.H); NCI Cancer Center Support Grant 2P30CA016520 (to J.T.); Linda and Paul Richardson Breast Cancer Research Funds (to J.T.); NIH NIGMS grant SC1GM121182 (to J.A.K.); ASCB MAC Visiting Professorship (to J.A.K.); and US-UK Fulbright-CRUK Scholar Award (to J.A.K.).

