## Supplemental Information for "A SNAI2-PEAK1 stromal axis drives progression and lapatinib resistance in HER2-positive breast cancer by supporting a cytokine expression profile that converges on PI3K/Akt signaling"

Sarkis Hamalian et al.

### FIGURES

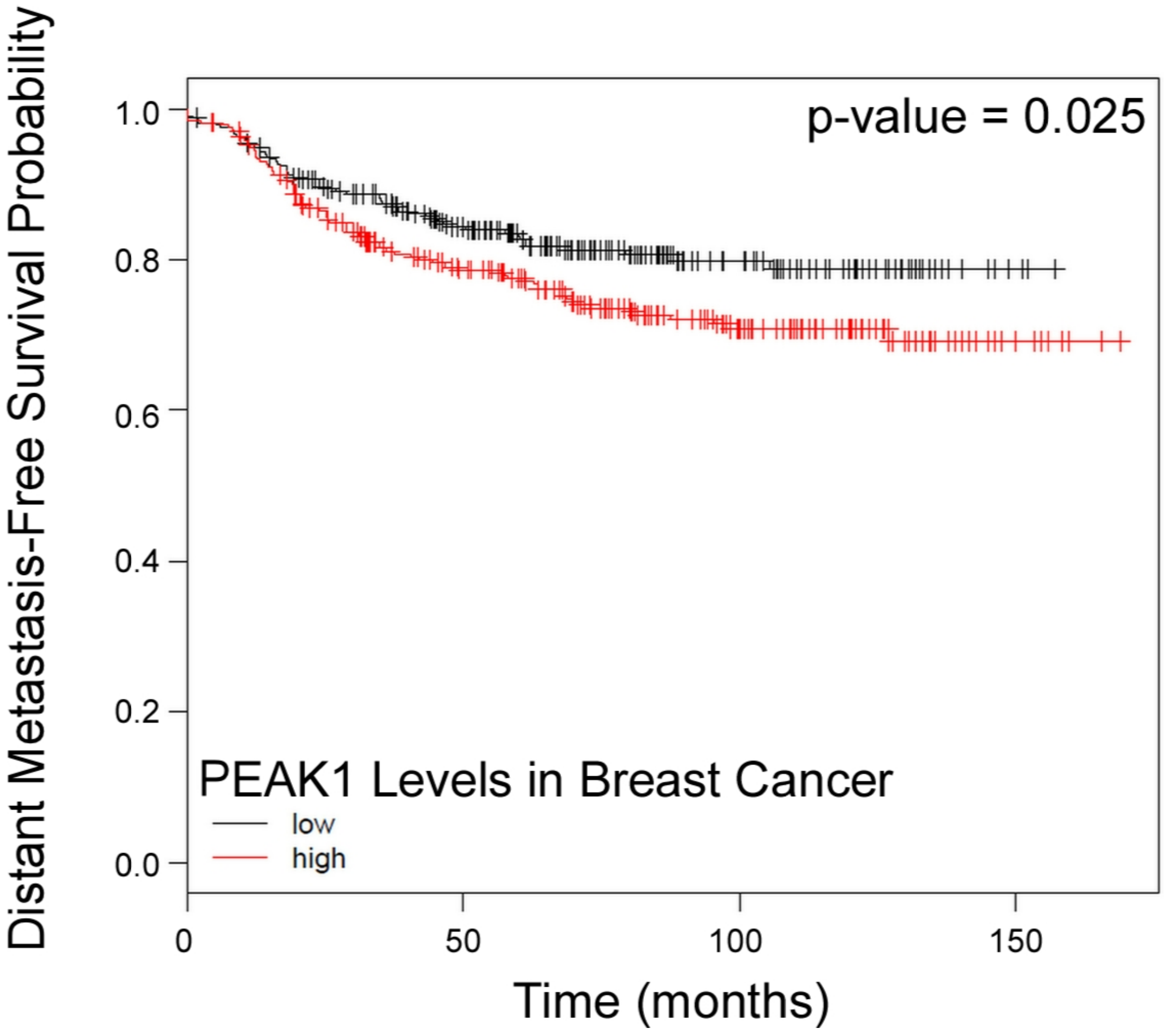


**Supplementary Figure 1:** Kaplan-Meier distant metastasis-free survival (DMFS) curves for low or high PEAK1 transcript levels in breast cancer patients (n = 1747).

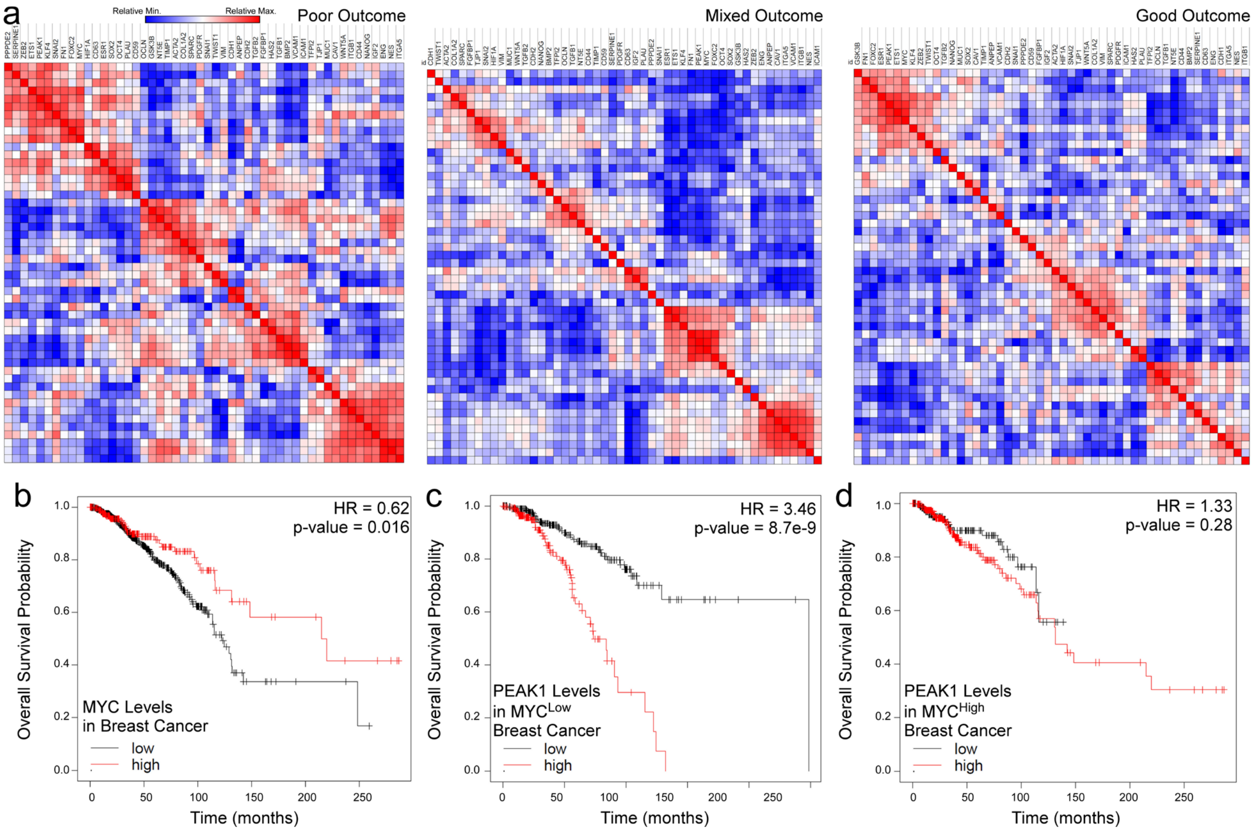


**Supplementary Figure 2:** a. Similarity matrix of average transcript levels for indicated genes in breast cancer stroma from patients categorized as having poor (left, n=8), mixed (center, n=27) or good (right, n=17) outcomes. b. Kaplan-Meier overall survival (OS) curves for patients with low or high MYC transcript levels across all breast cancer subtypes (n = 1089). c-d. Kaplan-Meier OS curves for low or high PEAK1 transcript levels in breast cancer patients selected for low (d) or high (e) MYC expression (n = 544).

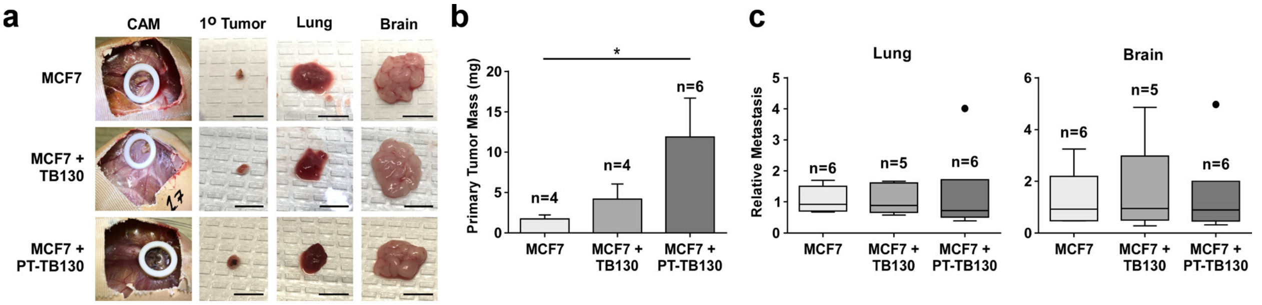


**Suplementary Figure 3:** a. Assay endpoint mages of indicated tissues for MCF7 cells, MCF7 cells + TB130 CAFs or MCF7 cells pre-treated (72 hours) with TB130 conditioned media and xenografted with TB130 CAFs (MCF7 + PT-TB130). b. Quantified primary tumor mass of experiment in (a). c. Relative metastasis of MCF7 cells in the lung (left) and brain (right) of experiment in (a). * indicates a p-value < 0.05 as determined by a One-Way ANOVA w/ multiple comparisons test.

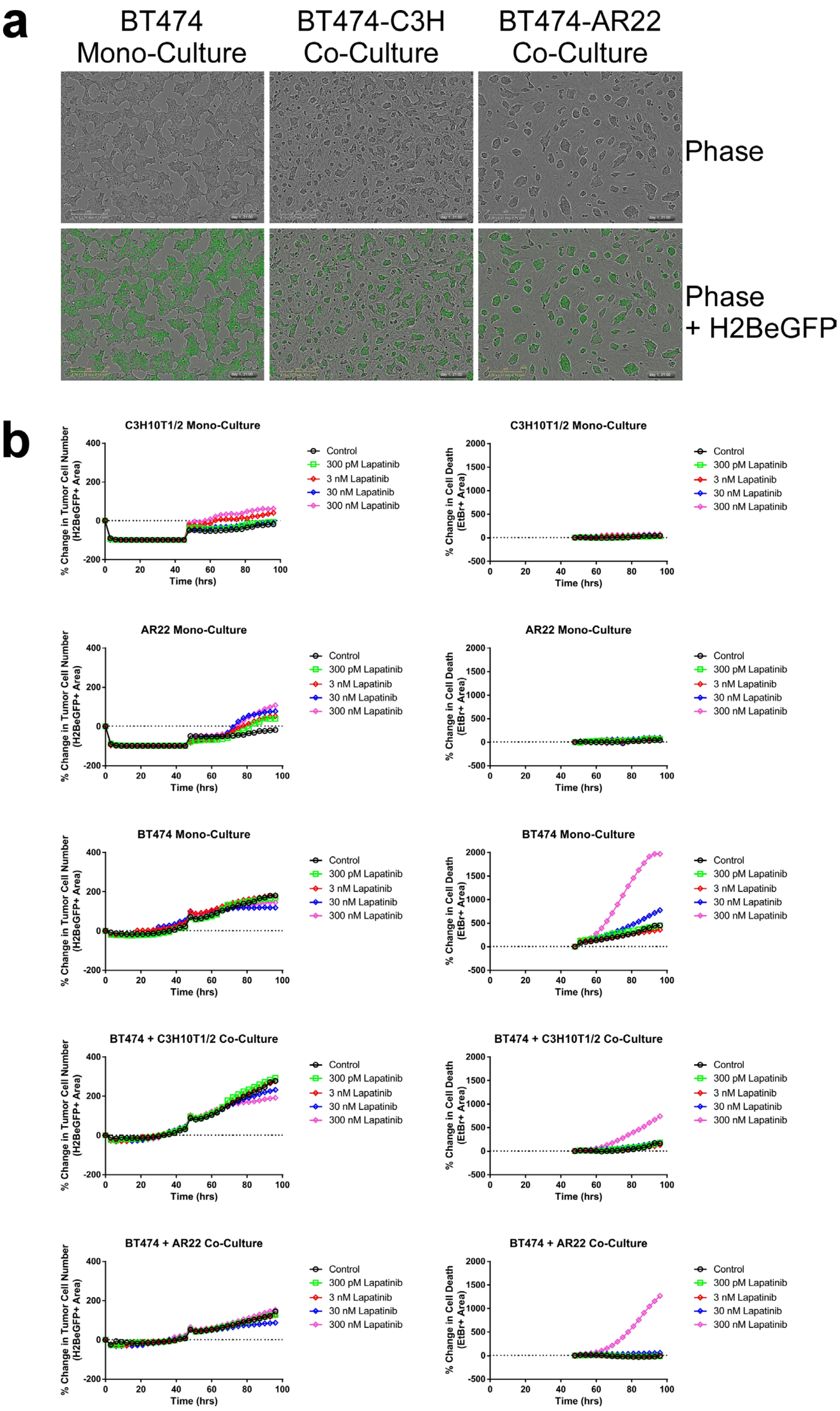


**Supplementary Figure 4:** a. Representative phase and phase+GFP overlay images for the indicated mono-culture or co-culture BT474 cells with either C3H10T1/2 MSCs or AR22 breast fibroblasts. b. Time-course traces for cell number measured by GFP signal (left) and cell death measured by EtBr signal (right) across all mono- or co-culture cell combinations and lapatinib treatment doses.

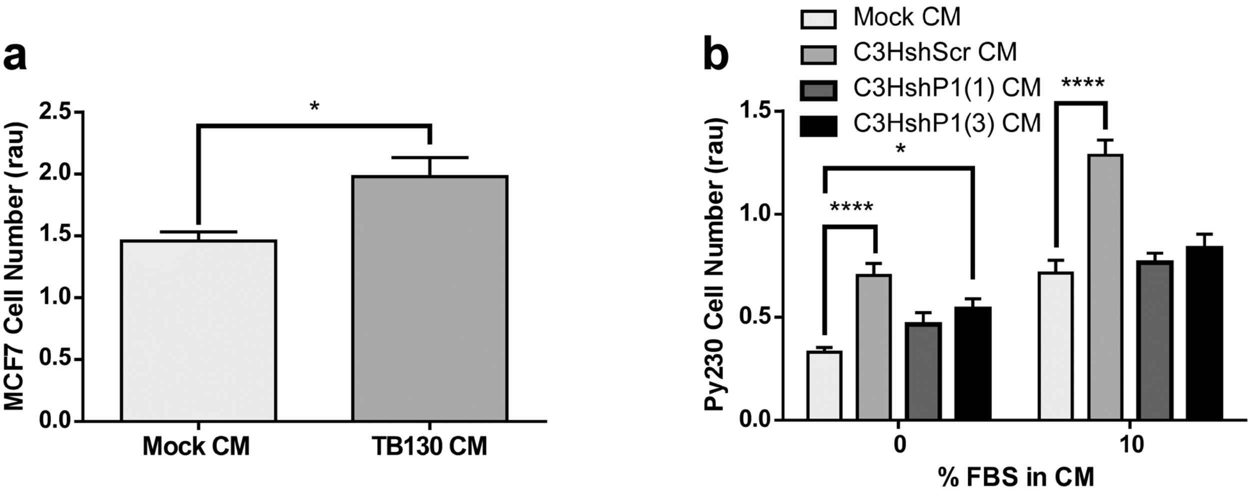


**Supplementary Figure 5:** a. Cell viability analysis of MCF7 cells treated with mock or TB130 CM. b. Cell viability analysis of mouse Py230 cells grown in serum-free or complete media and treated with mock CM or CM from the indicated shRNA derivatives of C3H10T1/2 cells. *, ***, or **** indicates a p-value < 0.05, 0.001 or 0.0001, respectively, as determined by a One-Way or Two-Way ANOVA w/ multiple comparisons post-test.

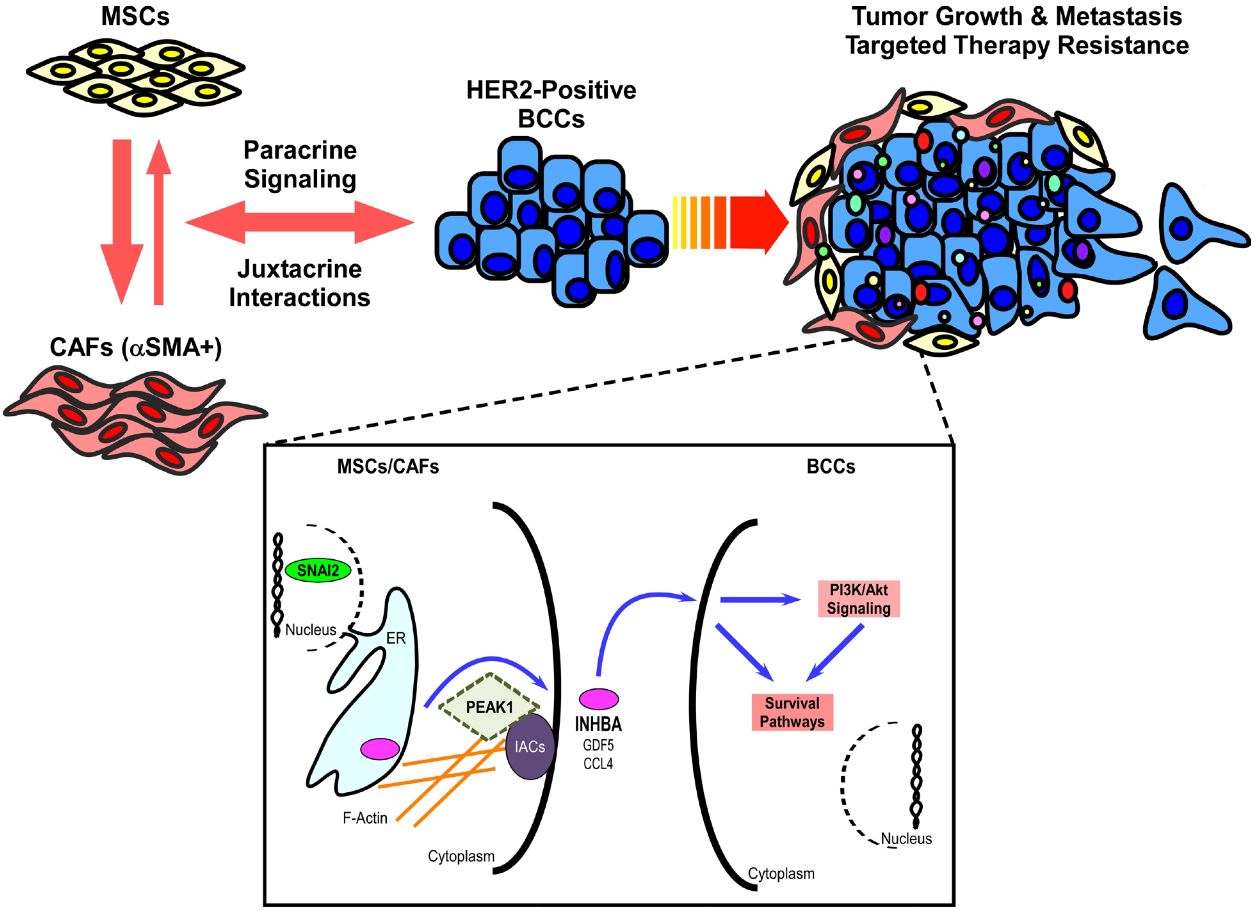


**Supplementary Figure 6:** Proposed model of mechanism by which stromal expression of PEAK1 drives tumor growth, metastasis and therapy resistance in neighboring HER2-positive breast cancer cells. Mesenchymal stem cells (MSCs) and/or cancer-associated fibroblasts (CAFs) interact with HER2-positive breast cancer cells (BCCs) in the primary tumor via aracrine and juxtacrine mechanisms. This favors conversion of MSCs to CAFs as indicated by their expression of markers such as alpha smooth muscle actin (αSMA). Inlay: PEAK1 and SNAI2 cooperate to promote poor outcomes in HER2-positive breast cancers enriched for MSC content. PEAK1, an actin cytoskeleton and integrin adhesion complex (IAC)-interacting protein is required for MSC expression of INHBA, GDF5 and CCL4. Increased levels of these activate the PI3K/Akt pathway in neighboring BCCs to further activate survival pathways and drive targeted therapy resistance.
